# Characterization of peroxidasin expression in histologically normal human adult and fetal kidney tissue

**DOI:** 10.1101/2023.12.21.572848

**Authors:** Isabel Brandão, Roberto Silva, Eduardo Conde, Bárbara Gomes, Paula Sampaio, Ana Costa Braga, Jorge Reis Almeida, Inês Soares Alencastre, João Paulo Oliveira

**Affiliations:** i3S – Instituto de Investigação e Inovação em Saúde, Universidade do Porto, Porto, Portugal; Institute of Biomedical Sciences Abel Salazar, Porto, Portugal; São João University Hospital Centre, Porto, Portugal; Faculty of Medicine, University of Porto, Porto, Portugal

**Keywords:** Peroxidasin, mature human kidney tissue, developing human kidney, compartment-specific analysis, distal tubules

## Abstract

Peroxidasin (PXDN) is an enzyme of the peroxidase family that plays a critical role in extracellular matrix (ECM) formation and tissue development. In the kidney, very few studies using animal cells suggested that PXDN may have a role in the maintenance of the structural and functional integrity of the renal ECM. Still, the normal expression and (patho)physiological roles of PDXN in the mature and developing human kidney remain unknown.

In this work we used fluorescent immunohistochemistry and advanced microscopy and image processing methodologies to perform a quantitative characterization of PXDN expression in the different anatomic compartments of histologically normal kidney tissue, obtained from adult nephrectomy specimens, and from postmortem fetal and neonatal kidney specimens (excluding death by kidney disease and genitourinary developmental anomalies).

Results show that PXDN is expressed in all mature kidney compartments, but at significantly higher levels in tubular epithelial cells, particularly in the distal tubules. A similar PXDN expression pattern was observed in the developing kidney, as early as the in the comma-shaped-body stage. In the fetal kidney, PXDN was diffusely expressed in the branches of the ureteric bud but not in the cells of the metanephric blastema.

This is the first demonstration that PXDN is differentially expressed in histologically normal human renal parenchyma, exhibiting a remarkably consistent pattern of predominant tubular expression since the early stages of metanephrogenesis This data suggests a relevant, compartment-specific, role of PXDN in nephrogenesis and in the kidney physiology, highlighting a main role in tubular functions, that is worth further investigation.

## Introduction

Peroxidasin (PXDN) is a multifunctional enzyme with a unique structure combining a catalytic peroxidase domain and several extracellular matrix (ECM) motifs, including leucine-rich repeats, immunoglobulin C2 (IgC2)-type domains, and a von Willebrand factor type C, that mediate protein-protein or protein-ligand interactions [24]. PXDN was originally identified 20 years ago, in the fruit fly *Drosophila melanogaster*, as an integral component of mature basement membranes (BMs) and a contributor to ECM consolidation, phagocytosis, and host defense [22]. Although the catalytic domain of the *Drosophila* PXDN showed homology with the human myeloperoxidase and eosinophil peroxidase, which are members of the same enzyme family, the human PXDN homolog was only identified in 2008 [9]. Since it exhibited highest tissue expression in the heart and the vascular wall, it was initially termed “vascular peroxidase 1” (VPO1). The demonstration that PXDN catalyzes the covalent crosslinking of type IV collagen (COL4) protomers directly within BMs, using hypohalous acid intermediates to form sulfilimine bonds (S=N) between a methionine sulfur and a lysine nitrogen [7]. shed new light on the biological role of the enzyme. Remarkably, the analysis of COL4 sequence homology evolution showed that the emergence of sulfilimine bonding, over 500 million years ago, coincided with the evolution of primitive BMs, implicating PXDN in a primordial innovation for tissue evolution, enabling tissues to withstand mechanical forces [7, 13]. Several lines of evidence have since implicated PXDN in tissue development [7, 13, 24, 29], endothelial cell survival and growth signaling [17], host defense [27], autoimmune disease pathogenesis [21], vascular remodeling [28, 34, 36], cardiovascular disease [20], diabetes [8], cancer [12, 37], and regulation of fibrosis [18, 23, 30].

Tissue development, homeostasis, and response to injury critically depend on cell-BM interactions, and perturbations of the macromolecular composition and stiffness of BMs, due to either injury or genetic defects, negatively affect cell-BM interactions. In the kidney, the BMs of glomeruli (GBM) and tubules (TBM) provide anchoring support respectively to podocytes and to the tubular epithelium; form permselective barriers limiting the bidirectional passage of macromolecules between the vascular space and the urinary space, in the glomeruli, and between the urinary filtrate and the interstitium, in the tubules; and are key dynamic mediators in embryonic development and organogenesis [1]. Therefore, defects in the architecture of kidney BMs are deleterious to cell-to-BM binding, which may result in loss of cell orientation and anomalous cell function, leading to tissue disruption; loss of GBM and TBM barrier functions; and to impaired tissue development and repair [1]. Indeed, genetic defects in the COL4 network, which forms the major structural scaffold of BMs, have a substantial impact on the mechanical stability of BMs, as observed in the GBM of patients with Alport syndrome [11]. The lower stiffness of kidney TBM observed in PXDN knockout mice, with deficient sulfilimination activity, provides direct experimental evidence that COL4 sulfilimine crosslinking contributes to the mechanical properties of kidney BMs, conferring structural reinforcement to the COL4 networks [6].

Yet, the only data suggesting the involvement of PXDN in the pathogenesis of kidney disease emerged from studies in the well-characterized transforming growth factor beta 1 (TGF-β1) signaling-dependent murine model of kidney fibrosis induced by unilateral ligation of the ureter [23]. In this model, PXDN was barely detectable in normal kidneys but its expression increased more than threefold after 7 days of ureter ligation, which was particularly enriched in the peritubular space. Notably, the colocalization of PXDN with fibronectin in the fibrotic kidneys resembled the *in vitro* findings of extracellular PXDN-containing fibrils partially colocalizing with fibronectin in cultures of TGF-β1-stimulated human pulmonary fibroblasts and human dermal fibroblasts [23]. Overall, the increased expression and its organization into a fibril-like network in the ECM observed in these experimental models, suggest that PXDN promotes matrix formation in response to injury, but whether the increased expression contributes to physiological or pathological fibrogenic responses is still unclear.

Although PXDN was found to be expressed in the human kidney by both mRNA profiling and proteome analyses [32], its differential distribution in the anatomic compartments of healthy human kidney is currently unknown. Herein, we provide the first description of the profile of PXDN expression in mature and developing histologically normal human renal parenchyma.

## Materials and Methods

*The used methodology is resumed in Figure 7 - Graphical abstract of the methodology workflow, in Supplementary Information*.

**Table 1.** Sources of histomorphologically normal mature and developing kidney tissue

|  |  |  |  |
| --- | --- | --- | --- |
| Mature kidney tissue – adult native and living-donor transplant nephrectomies |  |  |  |
| Reason for surgery | Cases | Gender | Age* |
| Renal cell carcinoma | 7 | M | median [range]: 50 y<br>[31-69] |
| Angiomyolipoma | 1 | F | 48 y |
| Blunt kidney trauma | 1 | M | 51 y |
| Early acute allograft renal artery thrombosis | 1 | M | 61 y |
| Developing kidney tissue – perinatal autopsy specimens |  |  |  |
| Fetal | 4 | M = 1 \| F = 3 | GA = 15, 17, 23, 30 w |
| Neonatal | 1 | F | GA<sub>B</sub> = 24 w / PPA = 3 w |
| * For nephrectomies: patients' ages at surgery, given in years (y); for perinatal deaths: gestational age (GA) or postpartum age (PPA) at the time of intrauterine or neonatal demise, given in weeks (w). GA<sub>B</sub> = GA at birth (w). M = male; F = female. |  |  |  |

### Retrieval and selection of mature and developing human kidney tissue specimens

Formalin-fixed paraffin-embedded (FFPE) specimens of mature (n = 10) and developing (n = 5) kidney tissue with normal histomorphological appearance and containing at least one whole glomerulus, were retrieved from the archives of the CHUSJ Pathology Department. Demographic and clinical details of the selected tissue sources are presented in Table 1. A margin of at least 1 cm from the tumor margins or the injured kidney parenchyma was a prerequisite for the collection of adult kidney specimens. Exclusion of developmental genitourinary anomalies and cause of death not imputable to kidney disease were conditions for preselecting the postmortem fetal and neonatal kidney specimens.

For subsequent use in this study, 3-μm-thick sections were cut from each of the FFPE blocks and mounted onto pseudonymized glass microscope slides, using the local routine laboratory procedures.

### Processing of the kidney tissue specimens for brightfield microscopy examination and manual segmentation

For brightfield microscopy examination, unstained as well as periodic acid–Schiff (PAS)-stained slides were prepared from sequential sections of each of the selected FFPE blocks, at room temperature (RT), according to standard histopathology techniques. In brief, the tissue sections were deparaffinized by washing in fresh xylene (3 x 10 min) and rehydrated to water by sequential exposure to decreasing concentrations of ethanol (1 x 10 min, at 100%, 96%, 70% and 50 %), before immersion in double-distilled water; for PAS staining, the tissue specimens were further oxidized in 0.5% periodic acid solution for 5 min, incubated with Schiff’s Reagent for 30 min, and counterstained in hematoxylin for 3 min. The unstained slides were used for manual segmentation of the kidney tissue compartments, and the PAS-stained slides as a topographical reference for the manual segmentation and for double-checking the normal histomorphology of the study specimens.

### Processing of the kidney tissue samples for indirect immunofluorescence analysis

After the deparaffinization and rehydration steps, the slides prepared for PXDN immunodetection were placed into sodium citrate antigen retrieval solution pH 6.0, heated for 20 min at 95ºC and left to cool at RT for an additional 20 min; washed with phosphate buffered saline (PBS); pH = 7.4); and incubated with blocking/permeabilization buffer (10% normal horse serum and 0.1% Triton X-100 in PBS) for 60 min at RT. Following blocking, the slides were incubated overnight at 4ºC with the primary antibody solutions diluted in blocking buffer; washed in PBS; and incubated for 1 h at RT with the secondary antibody solutions (see below), also diluted in blocking buffer. Finally, they were washed in PBS and incubated with 4′,6-diamidino-2-phenylindole (DAPI) solution in PBS, for cell nuclei counterstaining. After thorough washing with PBS, the slides were mounted with ProLong™ Gold Diamond Antifade Mountant (Thermo Fisher Scientific, Waltham, MA, USA; catalog # P36965), left to autopolymerize overnight at RT, and stored in the dark at 4ºC until analysis.

### Primary and secondary antibodies

For PXDN immunostaining, the primary antibody was a rabbit polyclonal anti-PXDN antibody (Abbexa; Cambridge, UK; catalog # abx101906), purified by antigen-specific affinity chromatography, and the fluorophore-conjugated secondary antibodies were either Donkey Anti-Rabbit IgG (H+L) Highly Cross-Adsorbed Secondary Antibody, Alexa Fluor™ Plus 647 (Invitrogen, Thermo Fisher Scientific; Waltham, MA, USA; catalog # A32795) or Abberior STAR 635P, Goat Anti-Rabbit IgG, 500µl (Abberior; Göttingen, German; catalog # ST635P-1002-500UG), respectively for standard immunofluorescence microscopy and for stimulated emission depletion (STED) high resolution microscopy.

To mark the proximal tubules, co-staining for Cluster of Differentiation 15 (CD15) antigen was carried out using the CONFIRM Anti-CD15 Mouse Monoclonal Primary Antibody (Ventana/Roche; Roche Diagnostics, Rotkreuz, Switzerland; catalog # 760-2504) and the fluorophore-conjugated Mouse IgG (H+L) Highly Cross-Adsorbed Secondary Antibody, Alexa Fluor™ Plus 568 (Invitrogen, Thermo Fisher Scientific; catalog # A10037).

The specificity of the commercial anti-PXDN primary antibody was checked by pre-adsorption testing. To this end, tissue fluorescence intensity was compared between slides that were stained with the primary antibody as provided by the manufacturer, or after the primary antibody was incubated with recombinant Human Peroxidasin Homolog Protein (Abbexa; catalog # abx068474), in equimolar (1:1) or 5x molar excess amounts. Incubation of slides with the secondary antibodies, omitting the primary antibody, was used as a negative control for the nonspecific binding of the secondary antibodies (Supplementary Information - Figure 6).

### Whole slide imaging, digital image analysis and virtual microscopy

The glass microscope slides processed for brightfield or fluorescence microscopy were converted into high-resolution digital images by high-speed scanning using a Nanozoomer S60 Digital Slide Scanner (Hamamatsu Photonics; Hamamatsu, Japan) or by a laser scanning confocal system equipped with White Light Laser, Leica Stellaris 8 stimulated emission depletion (STED) FAst Lifetime CONtrast (FALCON; Leica Microsystems; Wetzlar, Germany); super-resolution TauSTED images were additionally acquired with the Leica system using a 775 nm depletion laser and a 93x/1.3 glycerol objective. The adult kidney brightfield images were manually segmented into glomeruli, proximal and distal tubules, vessels, and interstitium (Figure 1), using the open source QuPath software (https://qupath.github.io/) [4]; fluorescence intensity measurements were obtained with ImageJ/Fiji [26] from the regions of interest defined on QuPath and after Rolling Ball Background Subtraction.

**Figure 1.**
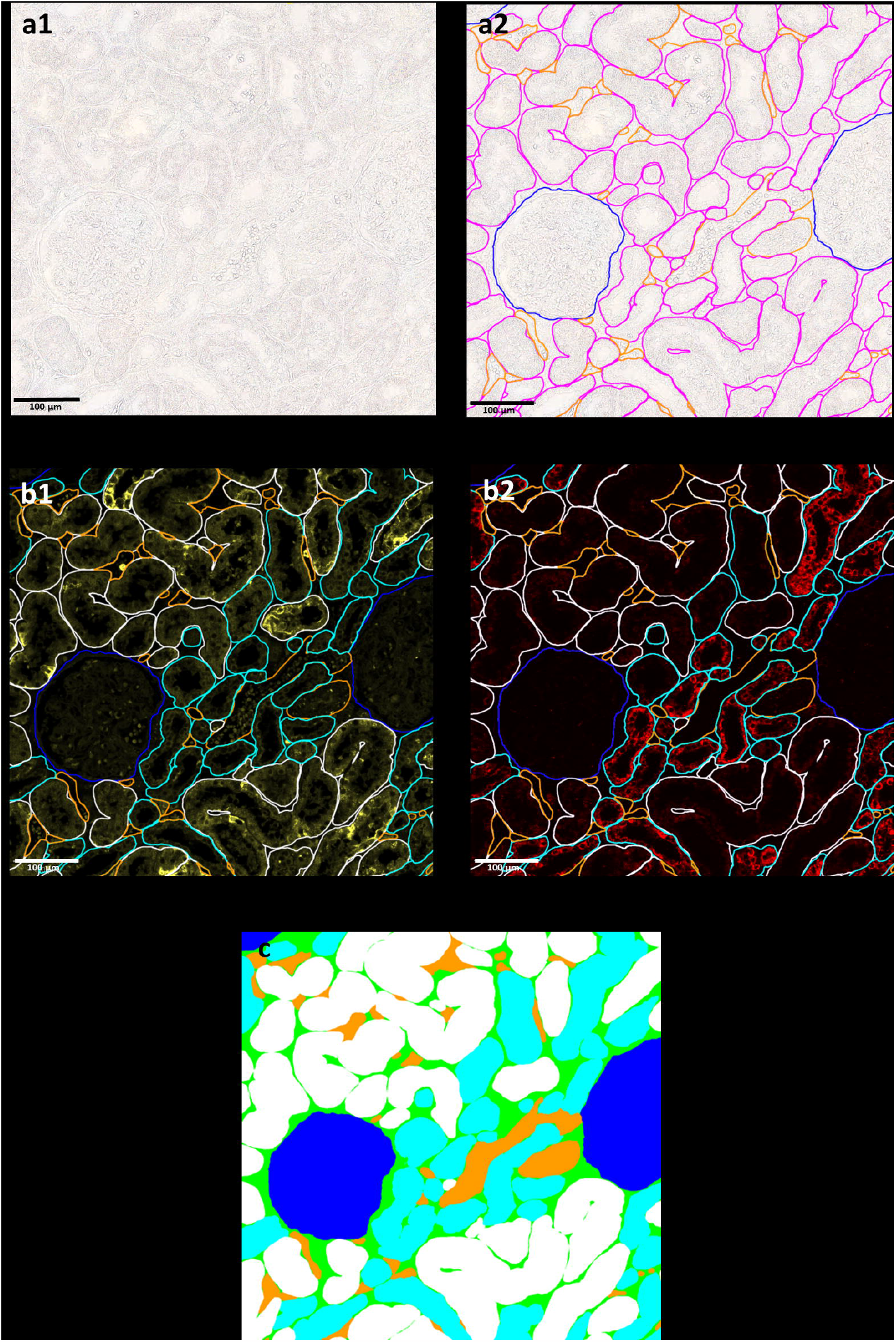
Workflow of image processing. After image acquisition [a1], the different kidney compartments were manually segmented in unstained brightfield images using the QuPath software [a2]. Annotations were subsequently transferred into CD15-labelled fluorescent images [b1] to distinguish the proximal tubules (CD15 positive) from the distal tubules (CD15 negative). The final annotations were then moved into peroxidasin (PXDN)-labelled fluorescent images [b2] and PXDN fluorescence intensity in the different kidney compartments was retrieved using ImageJ software. Filled map of annotations displaying the areas from where PXDN fluorescence intensity was acquired [c]. *Color codes ‒ glomeruli, deep blue; overall tubules, pink; distal tubules, turquoise; proximal tubules, white; vessels, orange; and interstitium, green*.

### Data analyses

Statistical analyses were performed with GraphPad Prism® (GraphPad Software; Boston, MA, USA) using one-way analysis of variance (ANOVA), followed by the post-hoc Tukey’s test for multiple comparisons where appropriate.

## Results

### Specificity of the immunofluorescent staining

Pre-adsorption of the primary anti-PXDN antibody with human recombinant PXDN led to a significant decrease of the intensity of the fluorescence signal in tested tissue samples, which was lost when the adsorption had been caried out with an excess of purified antigen (Supplementary Information – Figure 6). The absence of staining when the primary antibody was omitted from the staining protocol excluded nonspecific binding of the secondary antibodies (data not shown).

### PXDN expression in mature, adult kidney tissue

PXDN was expressed throughout the cortex and medulla (Figures 2 and 3), at comparable but relatively low levels across the glomerular, interstitial and vessel compartments, and at higher levels in the tubular compartment, particularly in the distal tubules. PXDN expression in the distal tubules was significantly higher than in the proximal tubules (p<0.0001), and it was also significantly higher in the tubular compartment than in the interstitium (p<0.0001 and p<0.05, respectively for the comparisons with the distal and the proximal tubules). In the glomerular, interstitial and vessel compartments, PXDN labeling was scarce, with a punctuate pattern of weak intensity, while in the tubules it was of higher intensity and diffuse, being visible in most of the cells included in the tubule cross-sections (Figure 2-a2 and 2-b2). No mesangial nor glomerular BM staining pattern was observed. Overall, PXDN staining was predominantly intracellular, exhibiting a distinct granular appearance in the cytoplasm that is particularly evident in the super resolution microscopy images (Figure 2-c3).

**Figure 2.**
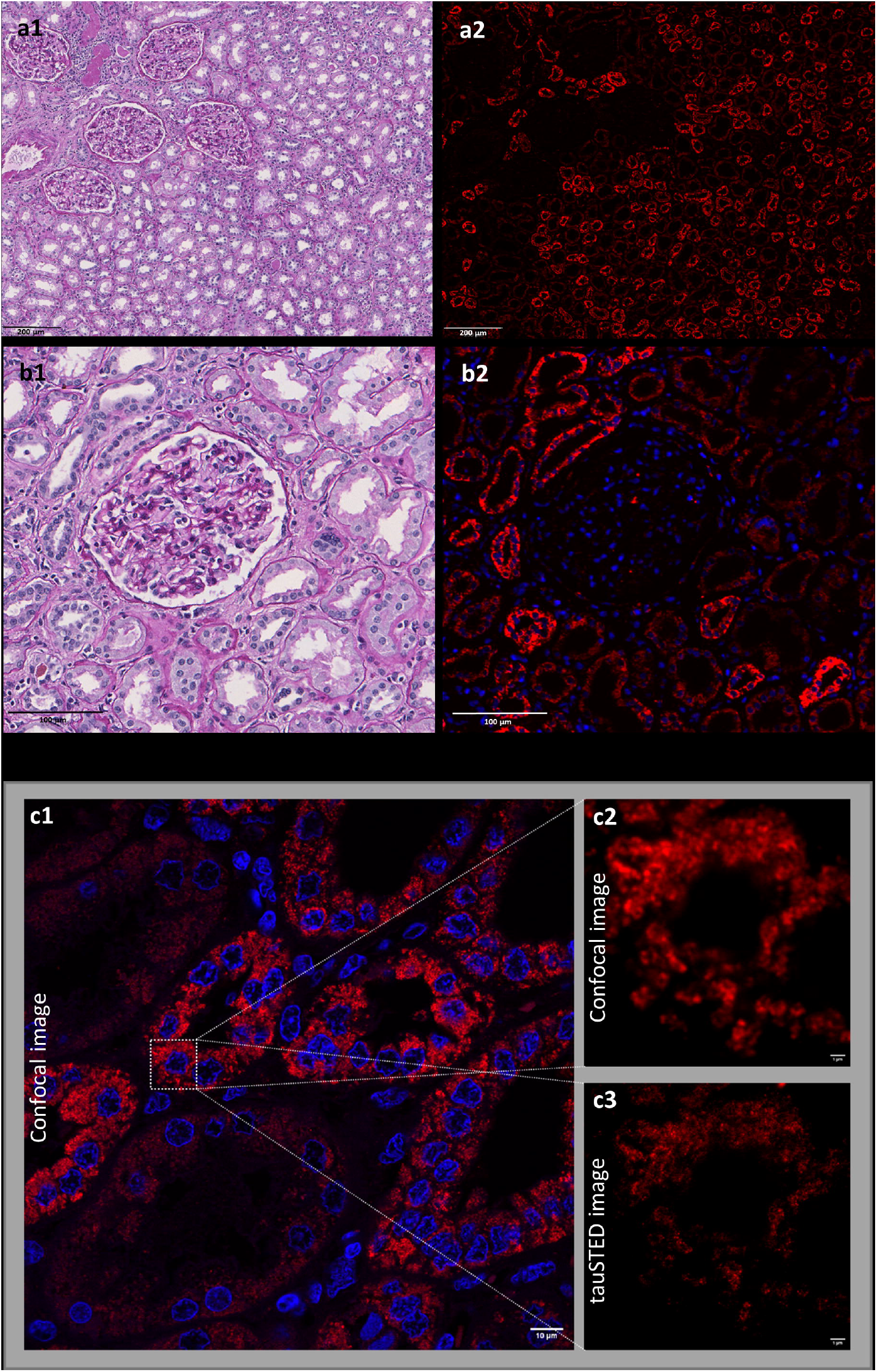
Exemplary micrographs of human kidney tissue samples from the histologically normal border of cancer nephrectomy specimens. Brightfield images of periodic acid–Schiff (PAS)-stained [a1, b1] and corresponding fluorescence images of 4′,6-diamidino-2-phenylindole (DAPI) and peroxidasin (PXDN) labeling [a2, b2, c] in the different kidney tissue compartments. Details of the histology and staining methods are provided in Materials and Methods. The secondary antibodies used for the immunofluorescence staining were either [a2, b2] the Donkey Anti-Rabbit IgG (H+L) Highly Cross-Adsorbed Secondary Antibody, Alexa Fluor™ Plus 647 (Invitrogen, Thermo Fisher Scientific; Waltham, MA, USA; catalog # A32795), or [c] the Abberior STAR 635P, Goat Anti-Rabbit IgG, 500µl (Abberior; Göttingen, German; catalog # ST635P-1002-500UG). No non-specific fluorescence labeling was observed in control samples when the secondary antibody was used in the absence of the primary rabbit anti-PXDN antibody (data not shown). The brightfield and fluorescence images were acquired in a NanoZoomer S60 Digital Slide Scanner (Hamamatsu Photonics; Hamamatsu, Japan) [a2, b2] or in a laser scanning confocal system equipped with White Light Laser, Leica Stellaris 8 STED FALCON (Leica Microsystems; Wetzlar, Germany) [c1, c2]; the super resolution TauSTED image [c3], allowing the observation of structures up to 50 nm resolution, was also acquired with the Leica system using a 775 nm depletion laser and a 93x/1.3 glycerol objective. Scale bars: [a] 200 µm, [b] 100 µm, [c] 10 µm and [c2, c3] 1 µm. *Color codes ‒ DAPI nuclei staining, blue; PXDN staining, red*.

**Figure 3.**
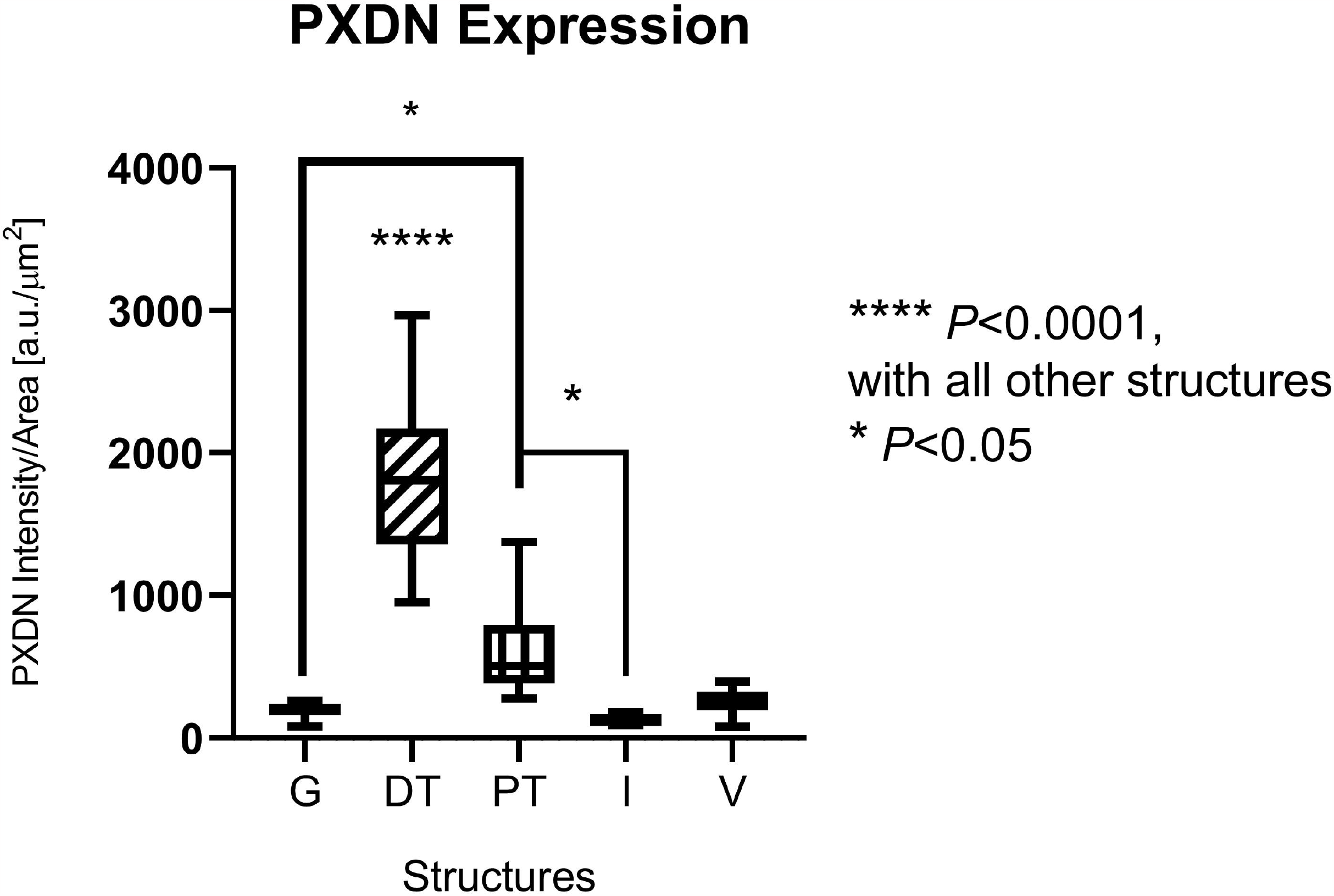
Quantitative analysis of peroxidasin (PXDN) expression in histologically normal adult kidney parenchyma according to kidney compartments. Statistical analysis was performed with GraphPad Prism® software using one-way ANOVA and Tukey’s multiple comparisons test (n = 10; median age = 50). Intensity/area values were obtained by dividing the integrated density by the total area of the compartment. *Abbreviations ‒ a*.*u*., *arbitrary units; G, glomeruli; DT, distal tubules; PT, proximal tubules; I, interstitium; V, vessels*.

### PXDN expression in the prenatal and postnatal, maturing kidney

Expression of PXDN in the undifferentiated metanephric mesenchyme was relatively scarce (Figure 4-a1), but it increased substantially in the condensed mesenchyme (Figure 4-a1), following induction by the branching ureteric bud, at the beginning of the nephrogenesis process [2], persisted at high levels through the comma-shaped and S-shaped stages, and at lower levels in the vascularized glomeruli. PXDN expression was conspicuous within the ureteric bud cells (Figure 4-a1), exhibiting a granular appearance along the basal side of the epithelium (Figure 4-b1, 4-b2). Unlike the mature glomeruli, PXDN was conspicuously expressed by both the cuboidal epithelial cells of Bowman’s capsule and the immature podocytes (Figure 4-c1, 4-c2) of the developing glomeruli, assuming a granular aspect in perinuclear location. There was no apparent expression of PXDN in the mesangial areas nor in the glomerular BMs.

**Figure 4.**
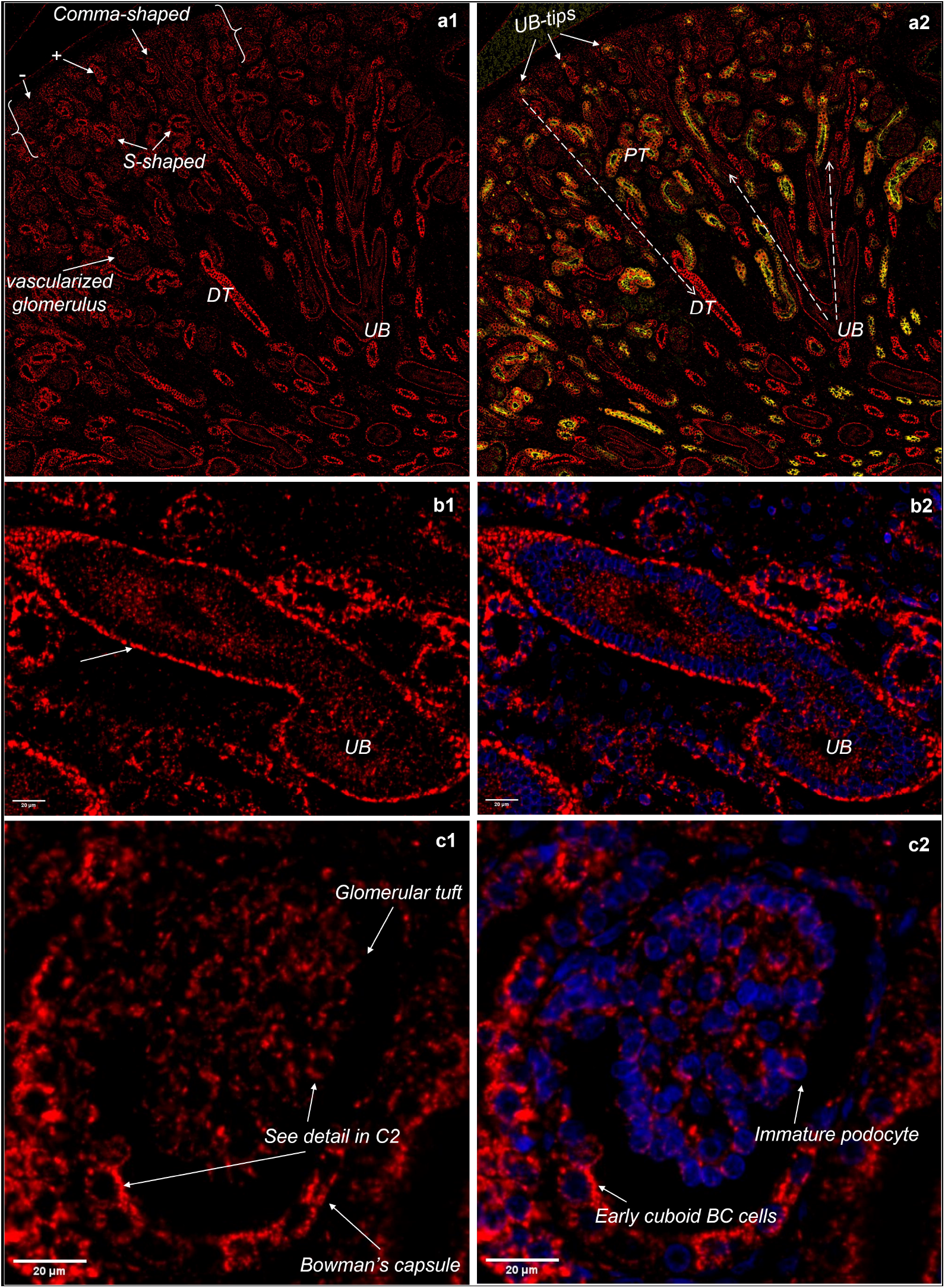
Exemplary fluorescence micrographs of peroxidasin (PXDN) expression in a 17-week-old fetus. [a1, a2] - The primordial nephrogenic zone ({…}), located at the periphery of the developing kidney, comprises areas of poorly cellular, undifferentiated metanephric mesenchyme (-), as well as areas with a large number of cells (+) undergoing mesenchymal-to-epithelial transition. This transition is triggered by reciprocal molecular interactions between the tips of the branching ureteric bud (UB) and adjacent metanephric mesenchymal cells, eventually leading to the formation of the renal vesicles, which are the most primitive nephron structures. Given the outward growth of the kidney, the morphological structures thar characterize the subsequent stages of nephrogenesis (comma-shaped body; S-shaped body; vascularized glomerulus) are located deeper within the developing kidney. Immunostaining for the Cluster of Differentiation 15 (CD15) antigen was used to distinguish proximal tubules (PT), which are CD15-positive, from distal tubules (DT), which do not express the CD15 antigen. [b1, b2] - Detail of a branching ureteric bud. PXDN is diffusely expressed by the UB epithelium, exhibiting a granular appearance at the basal side of the cells. [c1, c2] - Detail of a glomerulus with a well-developed capillary tuft. PXDN expression in the developing glomerulus is distinctly visible in the early cuboid cells of the Bowman’s capsule (BC) as well as in the immature podocytes, exhibiting a perinuclear granular appearance in both cell types. There is no significant expression of PXDN in the mesangial area nor in the glomerular basement membrane. Details of the histology and staining antibodies and methods are provided in Materials and Methods. *Abbreviations and color codes ‒ CD15 staining, yellow; DAPI, 4′*,*6-diamidino-2-phenylindole, cell nuclei staining, blue; PXDN, peroxidasin staining, red*.

Co-staining for PXDN and CD15 was observed at the tips of the ureteric bud and in the primitive and mature proximal tubules. In the distal tubules CD15 expression was absent while PXDN expression reached its maximum (Figure 4-a2). Similar relative expression patterns were observed in all maturing kidney tissue samples, irrespective of gestational age (data not shown).

The intensity of PXDN immunofluorescence during fetal development was quantitatively analyzed in 15, 17, 23, 27 and 30 weeks-old fetuses. In the ureteric bud (Figure 5-a), PXDN expression was detectable at relatively high levels as early as in the 15th week of gestation, declining progressively over the 3rd trimester of gestation. The number of ureteric buds identified in the 23-weeks-old fetal kidney was not high enough for statistical analysis and inclusion in the histogram. In the nephrogenic zone, the relative PXDN expression was two-fold higher in the differentiating as compared to the undifferentiated metanephric mesenchyme (Figure 5-b, 5-c). In both areas, the expression levels were highest in the 15-weeks-old fetal kidney, decreasing thereafter by about 50% and remaining relatively stable in the 3rd trimester. In the mature glomeruli, PXDN expression decreased progressively over the course of nephrogenesis, and even more in the adult kidney to about one third of the 3rd trimester levels (Figure 5-d). In the tubular compartment, PXDN expression decreased progressively over the course nephrogenesis and into the mature kidney, being significantly lower in the proximal tubules as compared to the distal tubules, in all stages of kidney development (Figure 5-e). In the distal tubules, the highest PXDN expression was in the 15-weeks-old fetal kidney, decreasing by more than 80% in the mature kidney (Figure 5-e)

**Figure 5.**
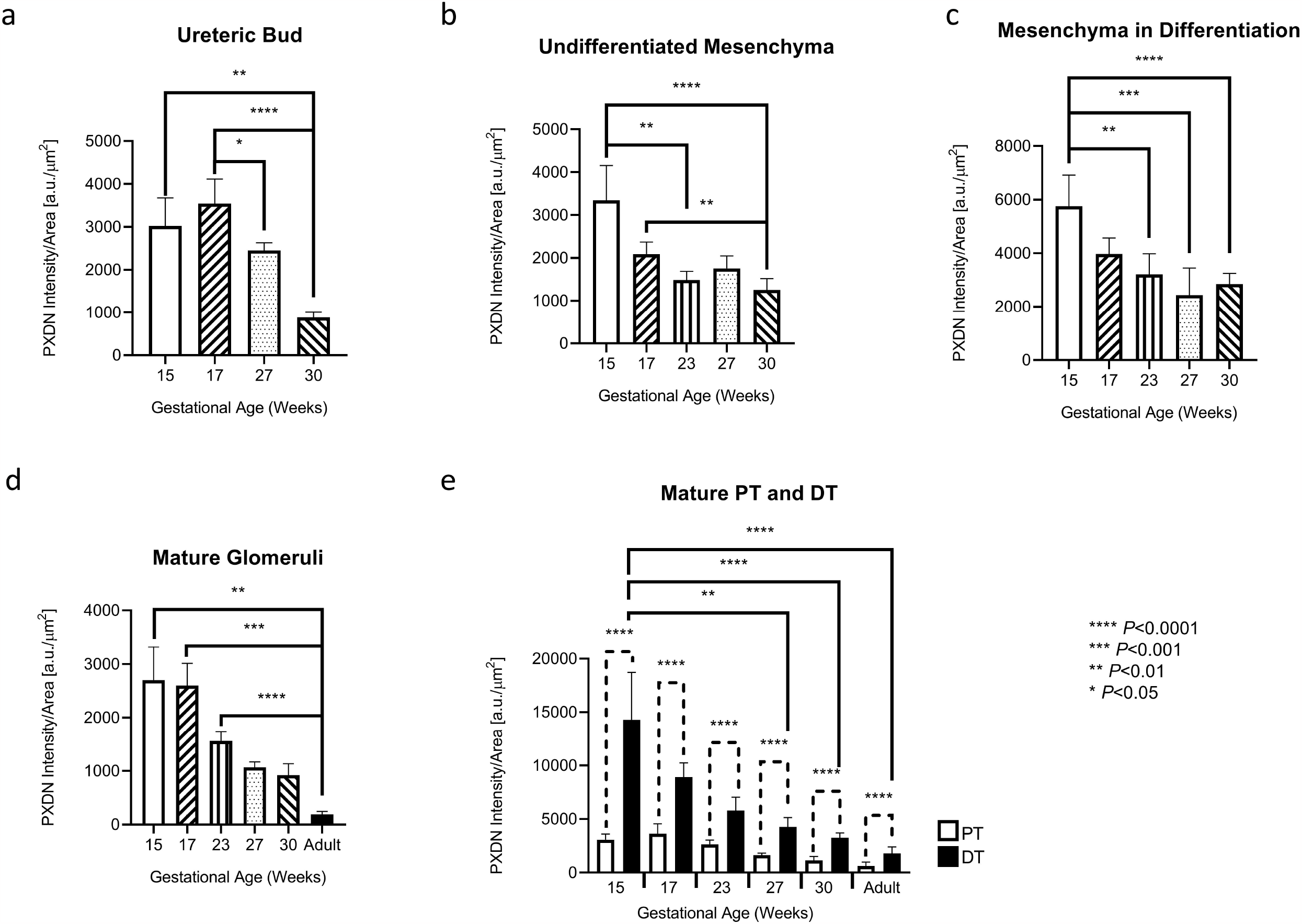
Comparative analyses of quantitative peroxidasin (PXDN) expression in developing kidneys at different gestational ages, and in mature kidneys. The quantification of PXDN expression in the areas comprising the ureteric bud[a], undifferentiated mesenchyme [b], mesenchyme in differentiation [c], mature glomeruli [d] and mature proximal (PT) and distal (DT) [e], was performed in kidney samples from 15, 17, 27 and 30 weeks-old fetuses. All statistical analyses were performed with GraphPad Prism® software using one-way ANOVA and Tukey’s multiple comparisons test. Results are displayed as the median with 95% confidence interval of 5-10 measurements in each kidney.

## Discussion

The different kidney tissue compartments have unique cellular constitutions, serving specific structural and functional purposes [3]. For that reason, and despite the highly laborious and time-consuming need for manual segmentation on virtual brightfield microscopy images, we chose to use a compartment-specific quantitative approach to describe the physiological expression of PXDN in the human kidney, along the different stages of nephrogenesis and in the mature organ. To the best of our knowledge these are entirely new data, providing a robust histological reference standard for studies addressing the role of PXDN in kidney development, in kidney (patho)physiology, or in the pathogenesis of chronic kidney disease, as well as those aiming to elucidate the genetic (dys)regulation of PXDN expression in such processes.

PXDN expression was identified in all mature kidney tissue compartments, but at significantly higher concentrations in the renal tubules, particularly the distal. The same expression pattern was observed in the developing kidney, as early as the induction of the nephrogenic blastema by the branching epithelial tips of the ureteric buds. Since PXDN is uniformly expressed by the ureteric bud epithelium, expression of PXDN by the condensed nephrogenic blastema seems to be related with the process of mesenchymal-to-epithelial transition. Notably, the highest expression of PXDN was observed in the distal tubules of a fetal kidney in the early 2^nd^ trimester of gestation. Overall, the physiological expression of PXDN in the human kidney is comparable to that described in the frog *Xenopus tropicalis*, where expression of Pxdn was already detectable in the primordium of the pronephric kidney, and persisted in the pronephric tubules and duct throughout development [31].

Taken together, our findings suggest that PXDN provides for essential roles in renal development and physiology, particularly in the tubular compartment. Considering the distinct metabolic requirements and the major energy pathways used by the proximal and the distal tubules [5, 25], the expression of PXDN in the tubular compartment may reflect its role in the regulation of mitochondrial membrane potential and oxidative stress pathways, as shown in prostate cancer cells [12].

The cytoplasmic granular appearance of the PXDN immunofluorescence signal in the kidney cells suggests that it predominantly localizes within subcellular organelles, most likely the endoplasmic reticulum, as demonstrated in human pulmonary and dermal fibroblasts [7]. Surprisingly, despite of the relevant physiological contribution of PXDN to the formation, architecture, and stability of BMs, and of its membrane binding motifs, neither the GBM nor the TBM stained for PXDN; this suggests that the biological role of PXDN in the BMs of the human kidney may occur mostly through transient, labile interactions, and that the protein is not incorporated, at an appreciable amount, into the BM structure.

Although the high degree of phylogenetic conservation of the *PXDN* gene [19] can be taken as evolutionary evidence of the importance of its biological role [35], pathogenic *PXDN* gene variants are associated with an autosomal recessive developmental disorder affecting the anterior segment of the eye [10, 15], without any clinically significant extraocular features. Furthermore, *Pxdn* knockout mice are born with an anterior segment eye defect replicating the corresponding human phenotype [6, 16, 33].

In conclusion, we have characterized the expression of PXDN in histologically normal developing and mature human kidney: PXDN was detected in all tissue compartments, although at significantly different degrees, reaching the highest levels in the distal tubular cells. This pervasive expression of PXDN in kidney tissue suggests a broader participation in physiological processes, beyond its well-known contribution to the biogenesis and stability of BMs. Further research into the elucidation of the normal functions of PXDN in the kidney, as well as into its likely role in key pathophysiological processes, particularly those involving redox-related ECM modifications or aberrant redox regulation [14], are worth the investment, offering the opportunity to discover new, potentially druggable pathways associated with progressive kidney disease.

## Statements and Declarations

### Ethics approval

This study was conducted in accordance with the Declaration of Helsinki. The research protocol was reviewed and approved by the institutional Health Ethics Commission (reference number CE-04-2023) and Data Protection Officer of *Centro Hospitalar Universitário de São João* (CHUSJ), Porto, Portugal.

### Authors contribution statements

I.S.A. and J.P.O. conceived, designed, and supervised the research project; I.B., J.R.A., and R.S. had major contributions to the experimental design; A.C.B., J.R.A. and R.S. collected and validated the kidney tissue specimens for study; B.G. provided histopathology technical support; I.B., I.S.A., J.R.A. and R.S. performed the acquisition for virtual microscopy; I.B. performed the manual segmentation of the kidney tissue compartments; I.B. did the statistical analyses; E.C. and P.S. provided expert assistance on virtual microscopy, fluorescence microscopy and digital image analysis; I.S.A. and J.P.O. wrote the draft manuscript. All authors discussed the results and read and approved the final manuscript. The corresponding author had full access to the data in the study and final responsibility for the decision to submit for publication.

## Acknowledgments

The kidney tissue samples used in this study were kindly provided by the Department of Anatomic Pathology of *Centro Hospitalar Universitário de São João* (CHUSJ), Porto, Portugal, which also granted training and access to the Nanozoomer S60 Digital Slide Scanner, from Hamamatsu Photonics.

## Supplementary Information

**Figure 6 – Competition assay to test anti-peroxidasin (PXDN) antibody specificity.**

Adult human kidney 3-µm-thick tissue sections were stained with rabbit polyclonal anti-PXDN antibody (Abbexa; Cambridge, UK; catalog # abx101906), in the absence [a] and presence of recombinant Human Peroxidasin Homolog Protein (Abbexa; catalog # abx068474), either in 1:1 equimolar [b] or 5x molar excess [c] amounts—in presence of the purified antigen, PXDN tissue staining was markedly reduced [b] or lost [c]. Fluorescence and brightfield images were acquired in NanoZoomer S60 Digital Slide Scanner (Hamamatsu Photonics; Hamamatsu, Japan). Scale bars: 20 µm. *Color codes ‒ 4′*,*6-diamidino-2-phenylindole (DAPI) nuclei staining, blue; PXDN staining, red*.

**Figure 7 – Graphical abstract of the methodology workflow**

## References

1. Abrahamson DR, Leardkamolkarn V (1991) Development of kidney tubular basement membranes Kidney Int 39:382–393. doi: 10.1038/ki.1991.50

2. Almeida JR, Mandarim-de-Lacerda CA (2002) Quantitative study of the comma-shaped body, S-shaped body and vascularized glomerulus in the second and third human gestational trimesters Early Hum Dev 69:1–13. doi: 10.1016/s0378-3782(02)00021-x

3. Balzer MS, Rohacs T, Susztak K (2022) How Many Cell Types Are in the Kidney and What Do They Do? Annu Rev Physiol 84:507–531. doi: 10.1146/annurev-physiol-052521-121841

4. Bankhead P, Loughrey MB, Fernandez JA, Dombrowski Y, McArt DG, Dunne PD, McQuaid S, Gray RT, Murray LJ, Coleman HG, James JA, Salto-Tellez M, Hamilton PW (2017) QuPath: Open source software for digital pathology image analysis Sci Rep 7:16878. doi: 10.1038/s41598-017-17204-5

5. Bhargava P, Schnellmann RG (2017) Mitochondrial energetics in the kidney Nat Rev Nephrol 13:629–646. doi: 10.1038/nrneph.2017.107

6. Bhave G, Colon S, Ferrell N (2017) The sulfilimine cross-link of collagen IV contributes to kidney tubular basement membrane stiffness Am J Physiol Renal Physiol 313:F596–F602. doi: 10.1152/ajprenal.00096.2017

7. Bhave G, Cummings CF, Vanacore RM, Kumagai-Cresse C, Ero-Tolliver IA, Rafi M, Kang JS, Pedchenko V, Fessler LI, Fessler JH, Hudson BG (2012) Peroxidasin forms sulfilimine chemical bonds using hypohalous acids in tissue genesis Nat Chem Biol 8:784–790. doi: 10.1038/nchembio.1038

8. Brown KL, Darris C, Rose KL, Sanchez OA, Madu H, Avance J, Brooks N, Zhang MZ, Fogo A, Harris R, Hudson BG, Voziyan P (2015) Hypohalous acids contribute to renal extracellular matrix damage in experimental diabetes Diabetes 64:2242–2253. doi: 10.2337/db14-1001

9. Cheng G, Salerno JC, Cao Z, Pagano PJ, Lambeth JD (2008) Identification and characterization of VPO1, a new animal heme-containing peroxidase Free Radic Biol Med 45:1682–1694. doi: 10.1016/j.freeradbiomed.2008.09.009

10. Choi A, Lao R, Ling-Fung Tang P, Wan E, Mayer W, Bardakjian T, Shaw GM, Kwok PY, Schneider A, Slavotinek A (2015) Novel mutations in PXDN cause microphthalmia and anterior segment dysgenesis Eur J Hum Genet 23:337–341. doi: 10.1038/ejhg.2014.119

11. Cosgrove D, Madison J (2022) Molecular and Cellular Mechanisms Underlying the Initiation and Progression of Alport Glomerular Pathology Front Med (Lausanne) 9:846152. doi: 10.3389/fmed.2022.846152

12. Dougan J, Hawsawi O, Burton LJ, Edwards G, Jones K, Zou J, Nagappan P, Wang G, Zhang Q, Danaher A, Bowen N, Hinton C, Odero-Marah VA (2019) Proteomics-Metabolomics Combined Approach Identifies Peroxidasin as a Protector against Metabolic and Oxidative Stress in Prostate Cancer Int J Mol Sci 20. doi: 10.3390/ijms20123046

13. Fidler AL, Vanacore RM, Chetyrkin SV, Pedchenko VK, Bhave G, Yin VP, Stothers CL, Rose KL, McDonald WH, Clark TA, Borza DB, Steele RE, Ivy MT, Aspirnauts Hudson JK, Hudson BG (2014) A unique covalent bond in basement membrane is a primordial innovation for tissue evolution Proc Natl Acad Sci U S A 111:331–336. doi: 10.1073/pnas.1318499111

14. Hanmer KL, Mavri-Damelin D (2018) Peroxidasin is a novel target of the redox-sensitive transcription factor Nrf2 Gene 674:104–114. doi: 10.1016/j.gene.2018.06.076

15. Khan K, Rudkin A, Parry DA, Burdon KP, McKibbin M, Logan CV, Abdelhamed ZI, Muecke JS, Fernandez-Fuentes N, Laurie KJ, Shires M, Fogarty R, Carr IM, Poulter JA, Morgan JE, Mohamed MD, Jafri H, Raashid Y, Meng N, Piseth H, Toomes C, Casson RJ, Taylor GR, Hammerton M, Sheridan E, Johnson CA, Inglehearn CF, Craig JE, Ali M (2011) Homozygous mutations in PXDN cause congenital cataract, corneal opacity, and developmental glaucoma Am J Hum Genet 89:464–473. doi: 10.1016/j.ajhg.2011.08.005

16. Kim HK, Ham KA, Lee SW, Choi HS, Kim HS, Kim HK, Shin HS, Seo KY, Cho Y, Nam KT, Kim IB, Joe YA (2019) Biallelic Deletion of Pxdn in Mice Leads to Anophthalmia and Severe Eye Malformation Int J Mol Sci 20. doi: 10.3390/ijms20246144

17. Lee SW, Kim HK, Naidansuren P, Ham KA, Choi HS, Ahn HY, Kim M, Kang DH, Kang SW, Joe YA (2020) Peroxidasin is essential for endothelial cell survival and growth signaling by sulfilimine crosslinkdependent matrix assembly FASEB J 34:10228–10241. doi: 10.1096/fj.201902899R

18. Liu Z, Xu Q, Yang Q, Cao J, Wu C, Peng H, Zhang X, Chen J, Cheng G, Wu Y, Shi R, Zhang G (2019) Vascular peroxidase 1 is a novel regulator of cardiac fibrosis after myocardial infarction Redox Biol 22:101151. doi: 10.1016/j.redox.2019.101151

19. Loughran NB, O’Connor B, O’Fagain C, O’Connell MJ (2008) The phylogeny of the mammalian heme peroxidases and the evolution of their diverse functions BMC Evol Biol 8:101. doi: 10.1186/1471-2148-8-101

20. Ma QL, Zhang GG, Peng J (2013) Vascular peroxidase 1: a novel enzyme in promoting oxidative stress in cardiovascular system Trends Cardiovasc Med 23:179–183. doi: 10.1016/j.tcm.2012.11.002

21. Manral P, Colon S, Bhave G, Zhao MH, Jain S, Borza DB (2019) Peroxidasin Is a Novel Target of Autoantibodies in Lupus Nephritis Kidney Int Rep 4:1004–1006. doi: 10.1016/j.ekir.2019.04.009

22. Nelson RE, Fessler LI, Takagi Y, Blumberg B, Keene DR, Olson PF, Parker CG, Fessler JH (1994) Peroxidasin: a novel enzyme-matrix protein of Drosophila development EMBO J 13:3438–3447. doi: 10.1002/j.1460-2075.1994.tb06649.x

23. Peterfi Z, Donko A, Orient A, Sum A, Prokai A, Molnar B, Vereb Z, Rajnavolgyi E, Kovacs KJ, Muller V, Szabo AJ, Geiszt M (2009) Peroxidasin is secreted and incorporated into the extracellular matrix of myofibroblasts and fibrotic kidney Am J Pathol 175:725–735. doi: 10.2353/ajpath.2009.080693

24. Peterfi Z, Geiszt M (2014) Peroxidasins: novel players in tissue genesis Trends Biochem Sci 39:305–307. doi: 10.1016/j.tibs.2014.05.005

25. Ratliff BB, Abdulmahdi W, Pawar R, Wolin MS (2016) Oxidant Mechanisms in Renal Injury and Disease Antioxid Redox Signal 25:119–146. doi: 10.1089/ars.2016.6665

26. Schindelin J, Arganda-Carreras I, Frise E, Kaynig V, Longair M, Pietzsch T, Preibisch S, Rueden C, Saalfeld S, Schmid B, Tinevez JY, White DJ, Hartenstein V, Eliceiri K, Tomancak P, Cardona A (2012) Fiji: an opensource platform for biological-image analysis Nat Methods 9:676–682. doi: 10.1038/nmeth.2019

27. Shi R, Cao Z, Li H, Graw J, Zhang G, Thannickal VJ, Cheng G (2018) Peroxidasin contributes to lung host defense by direct binding and killing of gram-negative bacteria PLoS Pathog 14:e1007026. doi: 10.1371/journal.ppat.1007026

28. Shi R, Hu C, Yuan Q, Yang T, Peng J, Li Y, Bai Y, Cao Z, Cheng G, Zhang G (2011) Involvement of vascular peroxidase 1 in angiotensin II-induced vascular smooth muscle cell proliferation Cardiovasc Res 91:27–36. doi: 10.1093/cvr/cvr042

29. Sirokmany G, Kovacs HA, Lazar E, Konya K, Donko A, Enyedi B, Grasberger H, Geiszt M (2018) Peroxidasin-mediated crosslinking of collagen IV is independent of NADPH oxidases Redox Biol 16:314–321. doi: 10.1016/j.redox.2018.03.009

30. Sojoodi M, Erstad DJ, Barrett SC, Salloum S, Zhu S, Qian T, Colon S, Gale EM, Jordan VC, Wang Y, Li S, Ataeinia B, Jalilifiroozinezhad S, Lanuti M, Zukerberg L, Caravan P, Hoshida Y, Chung RT, Bhave G, Lauer GM, Fuchs BC, Tanabe KK (2022) Peroxidasin Deficiency Re-programs Macrophages Toward Profibrolysis Function and Promotes Collagen Resolution in Liver Cell Mol Gastroenterol Hepatol 13:1483–1509. doi: 10.1016/j.jcmgh.2022.01.015

31. Tindall AJ, Pownall ME, Morris ID, Isaacs HV (2005) Xenopus tropicalis peroxidasin gene is expressed within the developing neural tube and pronephric kidney Dev Dyn 232:377–384. doi: 10.1002/dvdy.20226

32. Uhlen M, Fagerberg L, Hallstrom BM, Lindskog C, Oksvold P, Mardinoglu A, Sivertsson A, Kampf C, Sjostedt E, Asplund A, Olsson I, Edlund K, Lundberg E, Navani S, Szigyarto CA, Odeberg J, Djureinovic D, Takanen JO, Hober S, Alm T, Edqvist PH, Berling H, Tegel H, Mulder J, Rockberg J, Nilsson P, Schwenk JM, Hamsten M, von Feilitzen K, Forsberg M, Persson L, Johansson F, Zwahlen M, von Heijne G, Nielsen J, Ponten F (2015) Proteomics. Tissue-based map of the human proteome Science 347:1260419. doi: 10.1126/science.1260419

33. Yan X, Sabrautzki S, Horsch M, Fuchs H, Gailus-Durner V, Beckers J, Hrabe de Angelis M, Graw J (2014) Peroxidasin is essential for eye development in the mouse Hum Mol Genet 23:5597–5614. doi: 10.1093/hmg/ddu274

34. Yang L, Bai Y, Li N, Hu C, Peng J, Cheng G, Zhang G, Shi R (2013) Vascular VPO1 expression is related to the endothelial dysfunction in spontaneously hypertensive rats Biochem Biophys Res Commun 439:511–516. doi: 10.1016/j.bbrc.2013.09.012

35. Zerbino DR, Frankish A, Flicek P (2020) Progress, Challenges, and Surprises in Annotating the Human Genome Annu Rev Genomics Hum Genet 21:55–79. doi: 10.1146/annurev-genom-121119-083418

36. Zhang YS, He L, Liu B, Li NS, Luo XJ, Hu CP, Ma QL, Zhang GG, Li YJ, Peng J (2012) A novel pathway of NADPH oxidase/vascular peroxidase 1 in mediating oxidative injury following ischemia-reperfusion Basic Res Cardiol 107:266. doi: 10.1007/s00395-012-0266-4

37. Zheng YZ, Liang L (2018) High expression of PXDN is associated with poor prognosis and promotes proliferation, invasion as well as migration in ovarian cancer Ann Diagn Pathol 34:161–165. doi: 10.1016/j.anndiagpath.2018.03.002

