## Supplementary figures and images for "Characterization of peroxidasin expression in histologically normal human adult and fetal kidney tissue"

### Figure 6

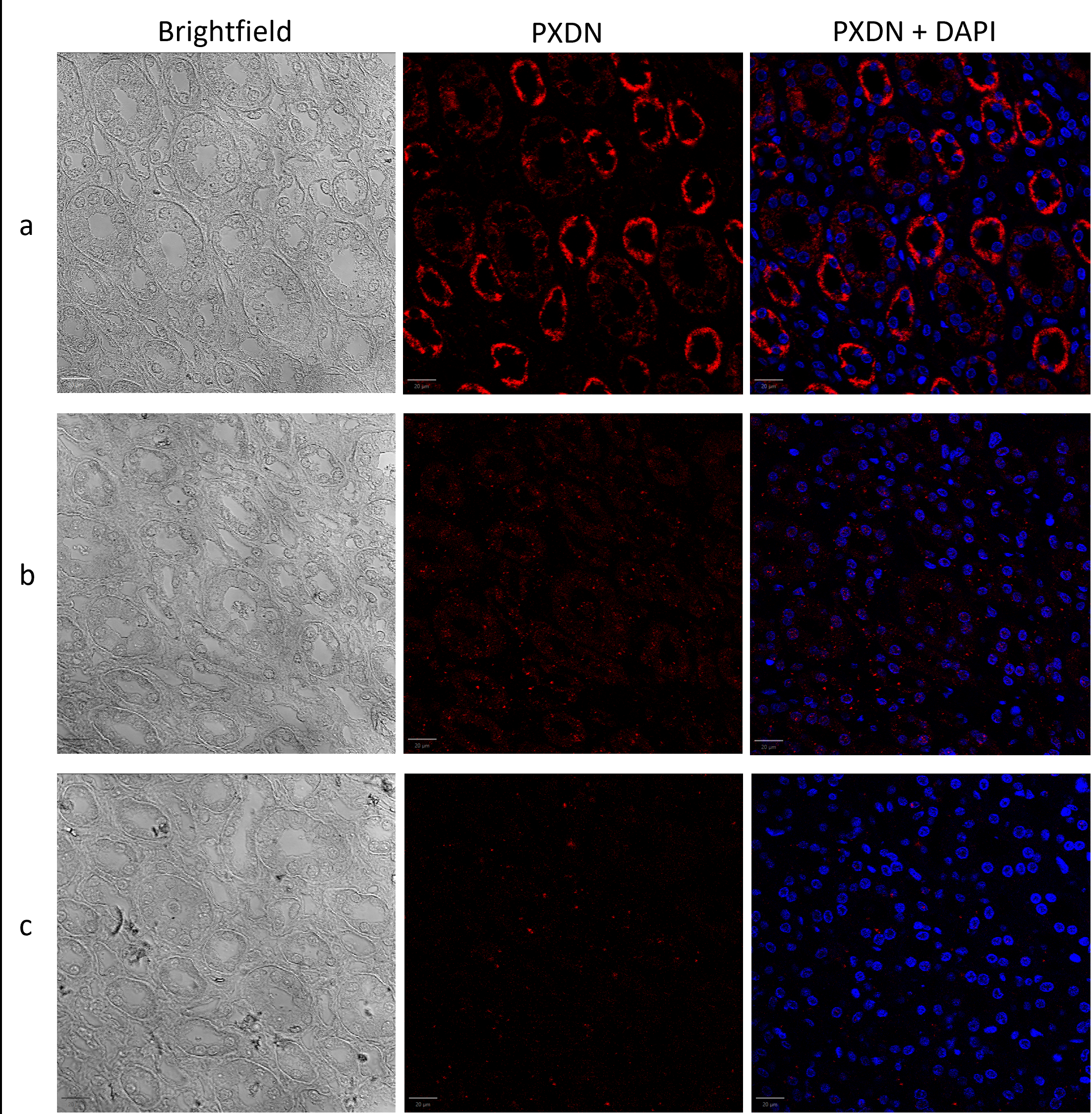

### Figure 7

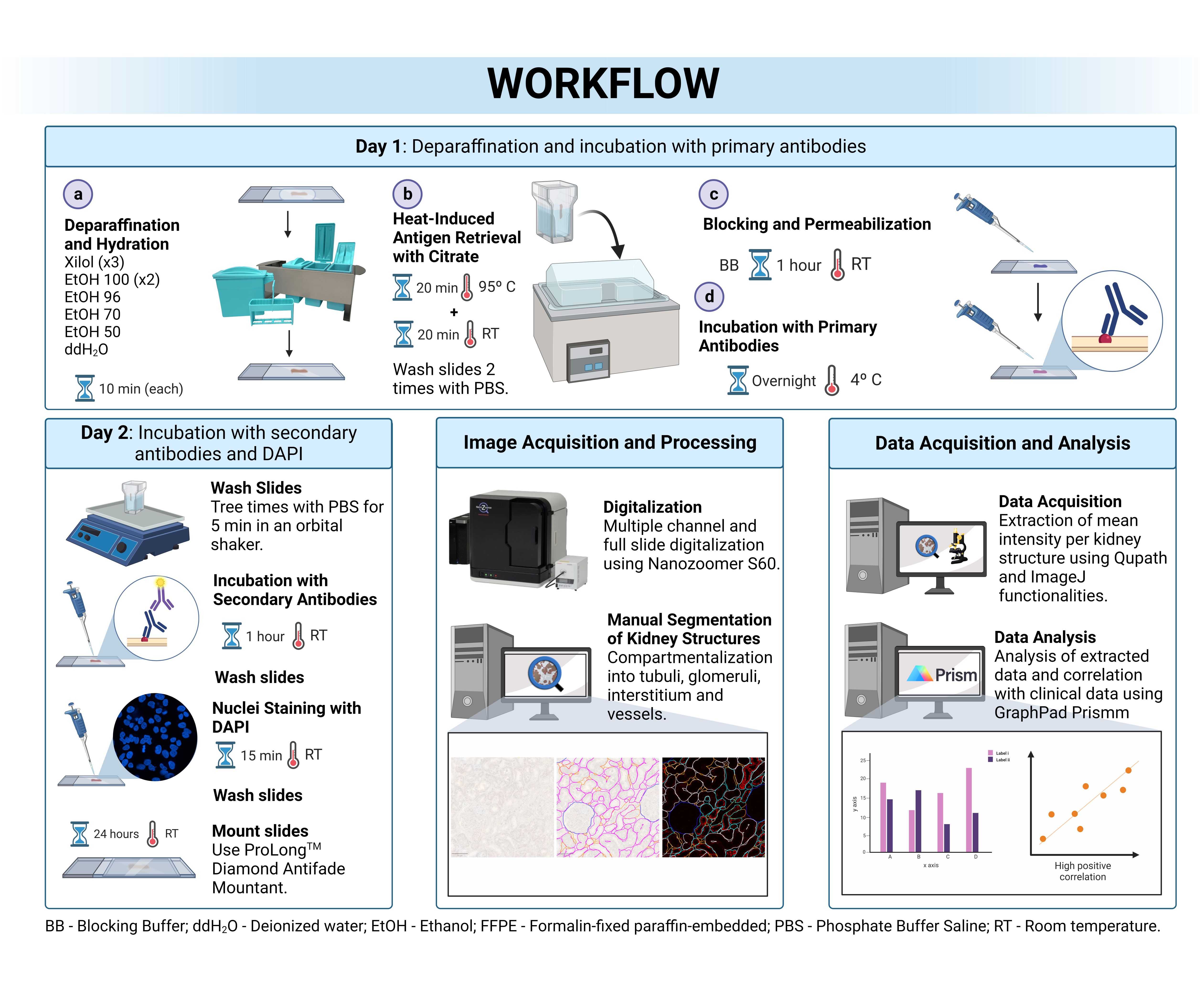
